# *Prevotella stercorea* links gut microbiome ecology to respiratory infection protection through a host-context-dependent, species-autonomous pathway

**DOI:** 10.64898/2026.05.25.727687

**Authors:** Ogochukwu Ofordile

## Abstract

Using a longitudinal cohort of 633 Gambian children (IHAT-GUT, NCT02941081), we resolve two mechanistically distinct ecological pathways linking *Prevotella stercorea* to infection risk. Its abundance positively predicts gut microbiome richness, consistent with community-level colonisation resistance for enteric outcomes. However, its association with reduced acute respiratory infection (ARI) persists unchanged after richness adjustment, identifying a species-autonomous pathway independent of community diversity. Weight-for-age z-score (WAZ) is uncorrelated with microbiome richness within strata, supporting WAZ as a proxy for host immune-metabolic reserve rather than a determinant of microbiome composition. In Low-WAZ children, *P. stercorea* at Day 1 associates with suppressed CRP, whereas in higher-WAZ children, elevated Day 1 inflammation predicts subsequent *P. stercorea* colonisation at Day 85, consistent with host-context-dependent immune selection. ARI and fever protection is richness-independent and concentrated in Low-WAZ children. *P. copri* does not retain an independent protective association when modelled jointly. These findings have direct implications for microbiome-directed interventions.

**Impact statement:** Gut *Prevotella stercorea* protects young children against respiratory infection through a species-specific, community-independent immune pathway that is most active in immunologically vulnerable hosts, defining a new mechanistic target for microbiome-directed respiratory therapeutics.

## Introduction

Species-level associations between the gut microbiome and infection are routinely reported, yet whether they reflect community properties such as diversity or direct species-autonomous effects is rarely tested. Recurrent infection in early childhood drives growth faltering and mortality in sub-Saharan Africa and South Asia (GBD 2019 Collaborators, 2020; Tickell et al., 2020) but the specific microbiome configurations conferring protection remain incompletely defined.

A central ecological contrast exists between industrialised and non-industrialised populations: *Bacteroides*-dominated microbiomes in the former, and *Prevotella*-dominated communities in the latter (Smits et al., 2017; De Filippo et al., 2010). Our prior longitudinal analysis of the IHAT-GUT cohort showed *Prevotella stercorea* was consistently depleted before infectious adverse events, ill children showed richness deficits, and *P. stercorea* formed part of an antagonistic network opposing an *Escherichia coli*-associated dysbiosis cluster (Ofordile et al., 2025). Microbiome composition preceded illness onset, establishing temporal ordering. We focus on *P. stercorea* and *P. copri* because they are the two dominant *Prevotella* species in this cohort, their co-occurrence enables within-genus specificity testing, and *P. stercorea* was identified as the primary infection-associated signal in our prior analysis.

Two competing frameworks generate testable predictions. In a community-mediated model, *Prevotella* contributes to microbiome diversity, with protection arising through richness-dependent colonisation resistance (Spragge et al., 2023). In a species-autonomous model, *P. stercorea* exerts effects independent of community structure, plausibly via gut–lung axis immune modulation (Dang and Marsland, 2019; Wypych et al., 2019). These are distinguished statistically: if mediated through richness, associations should attenuate after adjustment; if autonomous, they should persist. A third dimension - host immune context - defines the conditions under which each pathway is expressed, operationalised here as WAZ as a proxy for immune-metabolic reserve. WAZ is uncorrelated with microbiome richness within strata (Fig. 1), establishing that it modifies host response capacity rather than community composition. Interaction p-values and stratum-specific betas were adjusted using the Benjamini–Hochberg procedure within their respective comparison families; q-values are reported in Supplementary Tables 7 and 8A.

**Fig. 1.**
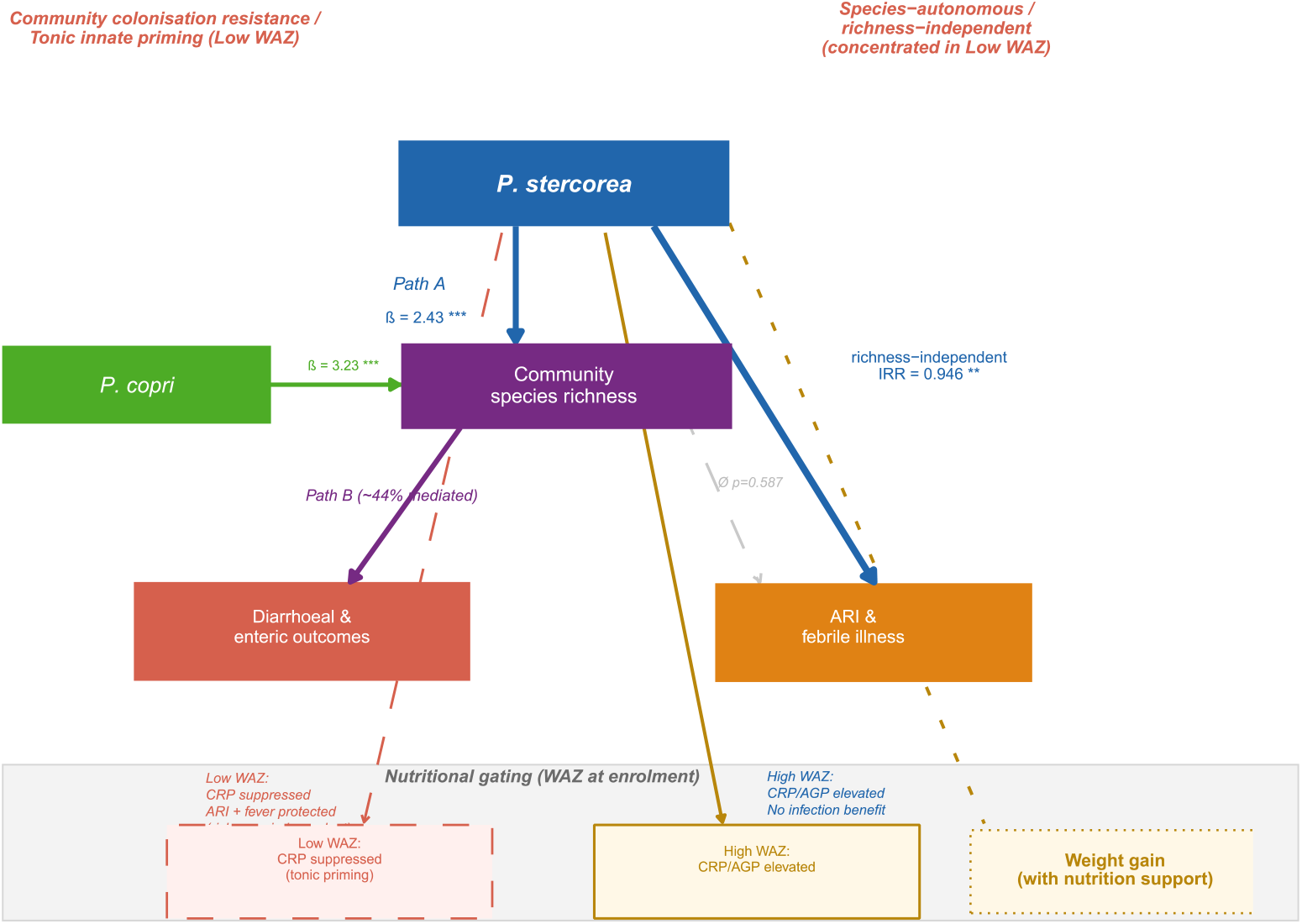
Conceptual framework: dual pathways with host immune context modulation. *P. stercorea* predicts community richness (Path A), contributing to colonisation resistance for enteric outcomes (left). Its association with ARI persists after richness adjustment, identifying a species-autonomous pathway (right), concentrated in Low-WAZ children where Day 1 colonisation associates with CRP suppression, consistent with tonic innate priming. In High-WAZ children, higher Day 1 inflammation predicts subsequent *P. stercorea* colonisation at D85 (immune-mediated colonisation selection) without infection benefit. WAZ is uncorrelated with richness within strata, confirming that host immune context operates through response capacity rather than community composition. Weight gain is predicted to require concurrent nutritional support (e.g. MDCF-2)(Chen et al., 2021; Mostafa et al., 2024).

Using longitudinal modelling and mediation analysis, we show that these frameworks are not mutually exclusive but compartment-specific. Enteric infection outcomes are consistent with community-mediated colonisation resistance (Foster and Bell, 2012), whereas respiratory infection outcomes are richness-independent and attributable specifically to *P. stercorea*, and not shared by the co-occurring *P. copri*. This species-resolved, compartmentalised framework links microbiome ecology to infection phenotypes and has direct implications for intervention design: whether protection is community-mediated or species-autonomous determines whether strategies should target community restoration or specific taxa.

## Results

### *Prevotella* species predict community richness

*P. stercorea* abundance was positively associated with species richness across all timepoints (D85: β = 2.43, 95% CI 1.77–3.09, p < 0.001; Table 1; Fig. 2), as was *P. copri* (D85: β = 3.23, 95% CI 1.97–4.48, p < 0.001), consistent across timepoints and confirmed in longitudinal mixed-effects models (Supplementary Table S1). The previously observed richness deficit in ill children (Ofordile et al., 2025) is therefore partly attributable to depletion of these taxa.

**Table 1.** *Prevotella* species abundance as predictors of community species richness.

| Species | Timepoint | n | $\beta$<br>(richness) | 95% CI | p |
| --- | --- | --- | --- | --- | --- |
| *P. stercorea* | D1 | 502 | 2.42 | 1.81–3.03 | < 0.001 |
| *P. stercorea* | D15 | 402 | 2.37 | 1.72–3.01 | < 0.001 |
| *P. stercorea* | D85 | 447 | 2.43 | 1.77–3.09 | < 0.001 |
| *P. copri* | D1 | 502 | 2.47 | 1.50–3.44 | < 0.001 |
| *P. copri* | D15 | 402 | 3.46 | 2.49–4.43 | < 0.001 |
| *P. copri* | D85 | 447 | 3.23 | 1.97–4.48 | < 0.001 |

OLS adjusted for age at sampling, sex, HAZ. $\beta$ = richness units per $\log(1 + \text{abundance})$ . BH-corrected q: all $p < 10^{-6}$ ; all $q < 0.001$ .

**Fig. 2.**
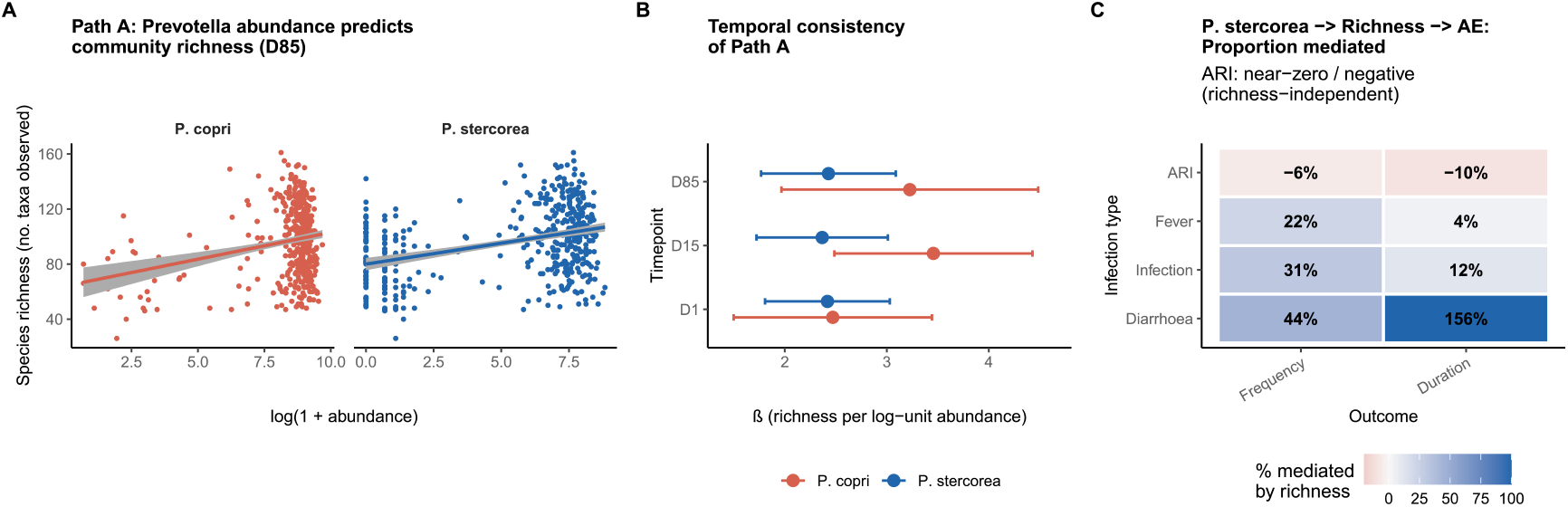
*P. stercorea* and *P. copri* predict gut microbiome richness. Mediation through richness by infection type. A) Richness vs log-abundance at D85 (n = 447), adjusted for age, sex, HAZ. (B) Effect estimates (β ± 95% CI) across D1, D15, D85, and mixed-effects model (C) Heatmap of proportion mediated (product-of-coefficients; n = 447). ARI: near-zero or negative. Diarrhoea: partial mediation. Colour scale −20% to 100%.

### Richness mediates enteric but not respiratory protection

Mediation analysis showed directionally consistent richness involvement in enteric outcomes: diarrhoea frequency showed 44% richness-mediated attenuation (path B β = −0.002, p = 0.183; Sobel p = 0.142), with diarrhoea duration providing corroborating directionality (Sobel p = 0.062). In contrast, ARI associations were not mediated by richness (−6% to −10%; Sobel p = 0.588, bootstrap ACME p = 0.532), richness was not associated with ARI (β = +0.002, 95% CI −0.005 to +0.009), and *P. stercorea* coefficients were unchanged after richness adjustment (Table 2; Fig. 2C; Supplementary Table S2). These findings support mechanistically distinct enteric and respiratory pathways.

**Table 2.**
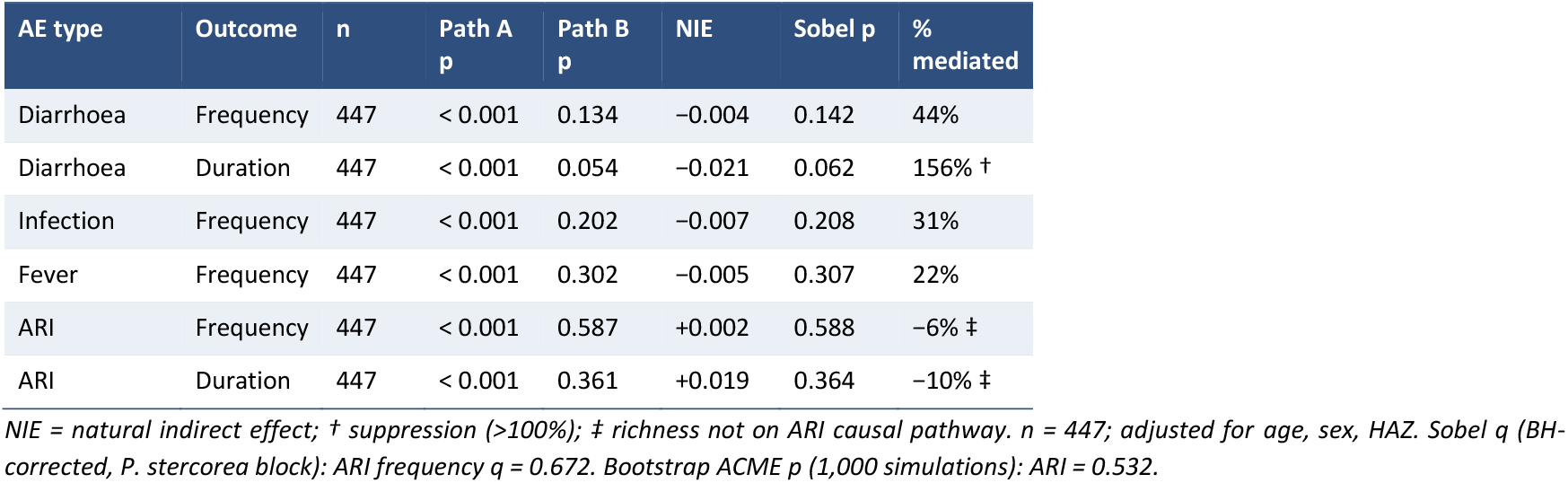
Proportion of *P. stercorea* association with adverse events mediated through species richness.

### *P. stercorea*–ARI association is species-specific and robust

*P. stercorea* associated with reduced ARI frequency (IRR = 0.946, 95% CI 0.913–0.981, p = 0.002, q = 0.009 after Benjamini–Hochberg correction), unchanged after richness adjustment (IRR = 0.942, 95% CI 0.907–0.979). In joint models with *P. copri* (r = 0.32), only *P. stercorea* retained significance; *P. copri* did not (β = −0.029, p = 0.381; Table 3; Fig. 3). Species specificity confirmed (joint model q: *P. stercorea* q = 0.016; *P. copri* q = 0.381). The association was not observed for diarrhoea or fever in the full cohort, further supporting compartment specificity.

**Table 3.**
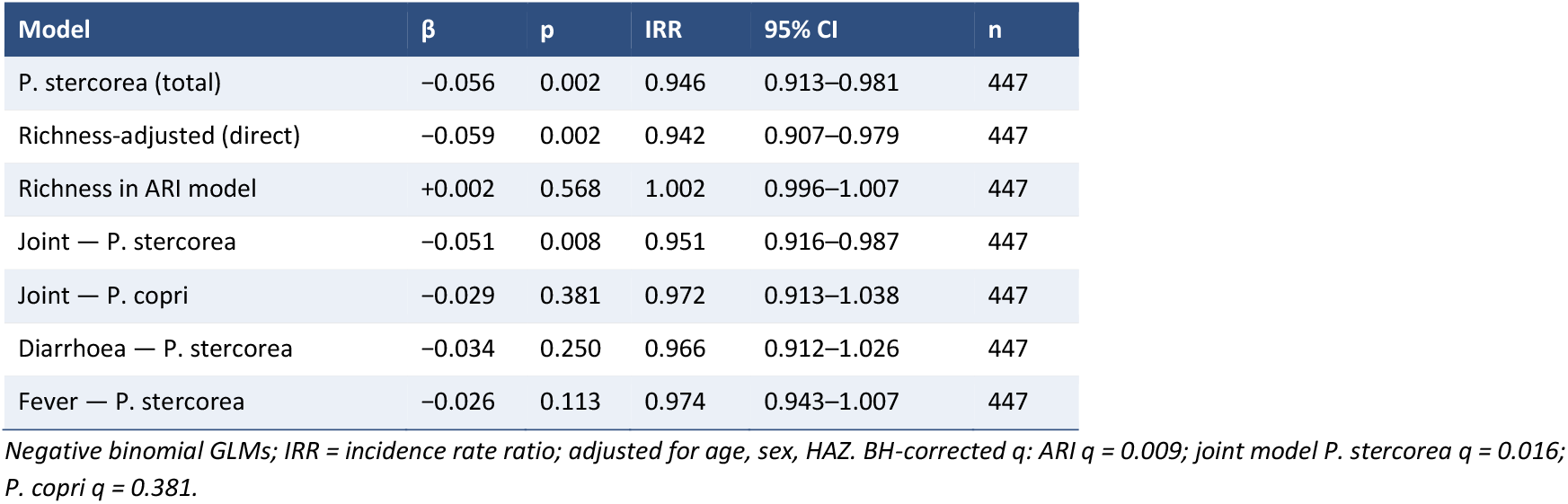
*P. stercorea* → ARI: total, richness-adjusted, and joint species models.

| Model | $\beta$ | p | IRR | 95% CI | n |
| --- | --- | --- | --- | --- | --- |
| *P. stercorea* (total) | −0.056 | 0.002 | 0.946 | 0.913–0.981 | 447 |
| Richness-adjusted (direct) | −0.059 | 0.002 | 0.942 | 0.907–0.979 | 447 |
| Richness in ARI model | +0.002 | 0.568 | 1.002 | 0.996–1.007 | 447 |
| Joint — *P. stercorea* | −0.051 | 0.008 | 0.951 | 0.916–0.987 | 447 |
| Joint — *P. copri* | −0.029 | 0.381 | 0.972 | 0.913–1.038 | 447 |
| Diarrhoea — *P. stercorea* | −0.034 | 0.250 | 0.966 | 0.912–1.026 | 447 |
| Fever — *P. stercorea* | −0.026 | 0.113 | 0.974 | 0.943–1.007 | 447 |

Negative binomial GLMs; IRR = incidence rate ratio; adjusted for age, sex, HAZ. BH-corrected $q$ : ARI $q = 0.009$ ; joint model *P. stercorea* $q = 0.016$ ; *P. copri* $q = 0.381$ .

**Fig. 3.**
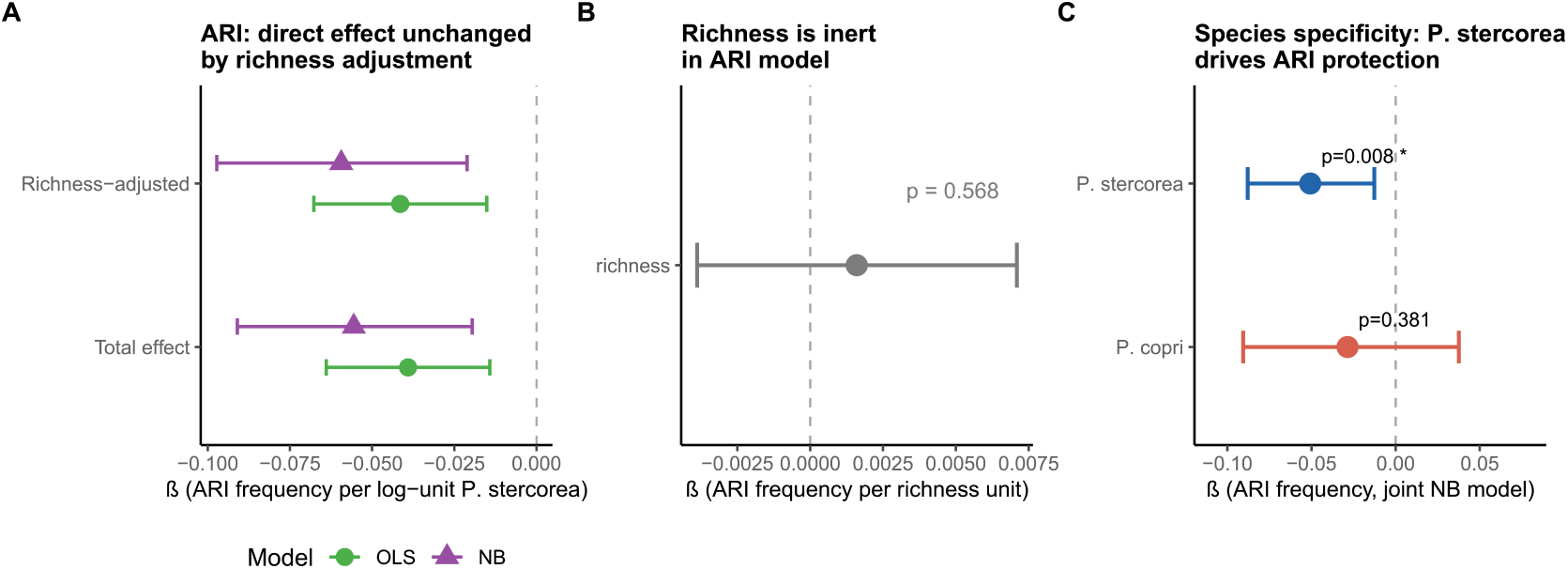
Species specificity of the ARI association. (A) ARI estimates before and after richness adjustment. (B) Richness in ARI model (non-significant). (C) Joint species model. Error bars: 95% CI.

### Host-context-dependent inflammatory phenotype and infection protection

In the full cohort, *P. stercorea* was not associated with CRP, calprotectin, or AGP, and baseline inflammation did not differ by illness status (all p > 0.5), arguing against pre-existing inflammation as a major confounder (Table 4; Supplementary Table S4).

**Table 4.** *P. stercorea* and inflammation biomarkers by WAZ stratum, and infection protection.

| Domain | Outcome / Marker | Timepoint | n | $\beta$ | 95% CI | p | Int. p | Sig |
| --- | --- | --- | --- | --- | --- | --- | --- | --- |
| **P.ste.–WAZ** | P.ste. vs WAZ (Low WAZ) | D85 | 248 | $r = +0.031$ | — | 0.627 | — | |
| | P.ste. vs WAZ (High WAZ) | D85 | 109 | $r = -0.079$ | — | 0.414 | — | |
| **WAZ–richness** | WAZ vs richness (Low WAZ) | D85 | 248 | $r = -0.016$ | — | 0.80 | — | |
| | WAZ vs richness (High WAZ) | D85 | 109 | $r = +0.081$ | — | 0.40 | — | |
| **Inflam. (D1→D1)** | CRP (Low WAZ) | D1→D1 | 276 | -0.037 | -0.067 to -0.007 | 0.016 | 0.015 | * |
|  | CRP (High WAZ) | D1→D1 | 121 | +0.007 | -0.043 to 0.057 | 0.783 |  |  |
|  | AGP (Low WAZ) | D1→D1 | 276 | -0.004 | -0.010 to 0.003 | 0.311 | 0.001 | ** |
|  | AGP (High WAZ) | D1→D1 | 121 | +0.014 | 0.004 to 0.025 | 0.008 |  | ** |
| **Inflam. (D85→D1)** | CRP (Low WAZ) | D85→D1 | 200 | -0.018 | -0.056 to 0.020 | 0.352 | < 0.001 | *** |
|  | CRP (High WAZ) | D85→D1 | 84 | +0.064 | 0.004 to 0.125 | 0.037 |  | * |
|  | AGP (Low WAZ) | D85→D1 | 200 | -0.007 | -0.015 to 0.001 | 0.076 | < 0.001 | *** |
|  | AGP (High WAZ) | D85→D1 | 84 | +0.018 | 0.005 to 0.031 | 0.007 |  | ** |
| **Infection protection** | ARI frequency (Low WAZ) | D85 exposure | 248 | -0.063 | -0.096 to -0.031 | < 0.001 | 0.012 | *** |
|  | ARI frequency (High WAZ) | D85 exposure | 109 | +0.008 | -0.036 to 0.052 | 0.712 |  |  |
|  | Fever frequency (Low WAZ) | D85 exposure | 248 | -0.058 | -0.095 to -0.021 | 0.003 | 0.048 | ** |
|  | Fever frequency (High WAZ) | D85 exposure | 109 | +0.012 | -0.034 to 0.058 | 0.607 |  |  |
| **Richness specificity (Low WAZ)** | Richness → ARI | D85 | 248 | -0.001 | -0.005 to 0.003 | 0.664 | — |  |
|  | Richness → Fever | D85 | 248 | -0.005 | -0.010 to 0.000 | 0.283 | — |  |
|  | ARI attenuation after richness adj. | D85 | 248 | < 12% | — | — | — |  |
| **Richness → Diarrhoea dur (Low WAZ)** | — | D85 | 248 | -0.015 | -0.026 to -0.004 | 0.008 | — | ** |
| **Richness → Diarrhoea dur (High WAZ)** | — | D85 | 109 | +0.004 | -0.015 to +0.023 | 0.699 | — |  |

OLS adjusted for HAZ, age, sex. D1→D1 = prospective; D85→D1 = reverse temporal. BH-corrected interaction $q$ (inflammation tests): CRP D1 $q = 0.018$ ; AGP D1 $q = 0.002$ ; CRP D85 $q = 0.002$ ; AGP D85 $q < 0.001$ . High WAZ stratified $q$ : AGP D1 $q = 0.020$ ; AGP D85 $q = 0.020$ ; CRP D85 $q = 0.063$ ( $n = 109$ ). Attenuation = $(c_{total} - c_{direct})/c_{total} \times 100$ . \* $p < 0.05$ ; \*\* $p < 0.01$ ; \*\*\* $p < 0.001$ .

After WAZ stratification, two temporally distinct and directionally asymmetric patterns emerged. First, in D1 stools → D1 biomarkers (prospective, same-timepoint), *P. stercorea* exhibited opposing associations by host immune context. In Low-WAZ children, Day 1 abundance associated with suppressed CRP (β = −0.037, int. q = 0.018), consistent with a lowered innate inflammatory set-point. In High-WAZ children, *P. stercorea* associated with elevated AGP (β = +0.014, int. q = 0.002), consistent with immune engagement under higher metabolic reserve.

Second, in D85 stools → D1 biomarkers (reverse temporal direction), associations were confined to High-WAZ children: higher baseline CRP and AGP predicted higher subsequent *P. stercorea* at D85 (CRP: β = +0.064, p = 0.037, int. q = 0.002; AGP: β = +0.018, p = 0.007, int. q < 0.001), whereas no associations were observed in Low-WAZ children. CRP High WAZ stratified q = 0.063 (n = 109; directionally consistent with AGP pattern, q = 0.020). This reverse-temporal pattern indicates that baseline inflammatory tone predicts subsequent colonisation in higher-immune-reserve children, rather than *P. stercorea* driving inflammation, defining a host-context-dependent, directionally asymmetric coupling between inflammatory state and colonisation.

Infection protection followed the same host-context stratification. In Low-WAZ children, *P. stercorea* associated with reduced ARI frequency (β = −0.063, p < 0.001), ARI duration (β = −0.282, p < 0.001), fever frequency (β = −0.058, p = 0.003), and fever duration (β = −0.287, p = 0.001), with significant WAZ interactions (ARI frequency p_interaction = 0.012; fever frequency p_interaction = 0.048; Supplementary Table S7). No associations were observed in High-WAZ children. Richness also did not predict diarrhoea outcomes in High-WAZ children (frequency p = 0.644; duration p = 0.699), indicating that the richness-mediated enteric protection observed in Low-WAZ children does not extend to better-nourished children, and that neither mechanistic pathway confers detectable protection in this stratum.

These effects were independent of community diversity. Within Low-WAZ children, richness was not associated with ARI (p = 0.664) or fever (p = 0.283), and *P. stercorea* effect estimates were minimally attenuated after richness adjustment (<12%), supporting a species-autonomous pathway. In contrast, diarrhoea frequency showed 44% richness-mediated attenuation in the full cohort, with directionally consistent signal from diarrhoea duration, supporting community-mediated enteric protection. *P. copri* showed no significant WAZ interaction for any inflammation outcome (all interaction q > 0.13 after BH correction), indicating that host-context-dependent immune modulation is not a shared genus-level property but is specific to

Notably, the Low-WAZ stratum in which Day 1 CRP is suppressed is the same stratum in which richness-independent protection against ARI and fever is concentrated (Fig. 4; Table 4; Supplementary Table S8), indicating mechanistic co-localisation of inflammatory modulation and clinical protection.

**Fig. 4.**
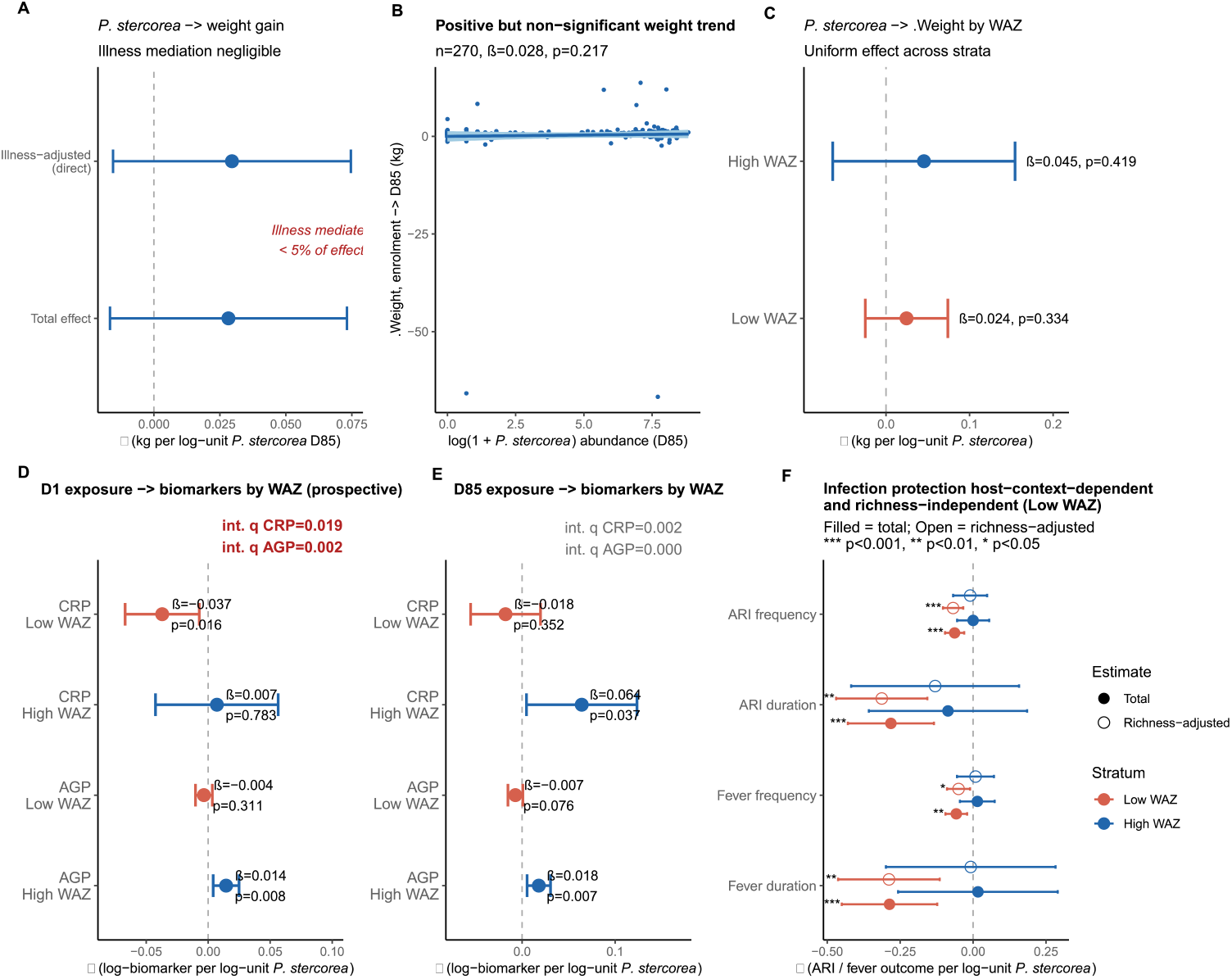
*P. stercorea*: weight gain, biomarker stratification, and mechanistic co-localisation. (A) Weight gain estimates with and without illness adjustment; < 2% mediated. (B) Scatter plot (n = 270). (C) WAZ-stratified weight gain estimates; flat interaction (pinteraction = 0.143). (D) D1→D1 biomarker associations by WAZ (prospective; primary host-context test). Low WAZ: CRP suppression (β = −0.037, int. q = 0.018). High WAZ: AGP elevation (β = +0.014, int. q = 0.002). ΔWeight int. q = 0.143. Full q-values in Table 4. (E) D85→D1 (reverse temporal): High-WAZ children with higher Day 1 CRP/AGP subsequently have higher *P. stercorea* at D85 (both interactions p < 0.001); null in Low WAZ. Interpreted as inflammatory tone predicting colonisation, not *P. stercorea* driving inflammation. (F) WAZ-stratified infection protection: *P. stercorea* → ARI and fever concentrated in Low WAZ (Low WAZ q ≤ 0.006); absent in High WAZ. Filled = total effect; open = richness-adjusted direct effect; attenuation < 12%. Dashed line at zero.

### WAZ is independent of richness and *P. stercorea* abundance within strata

Within WAZ strata, richness was not associated with WAZ (Low WAZ: r = −0.016, p = 0.80; High WAZ: r = +0.081, p = 0.40), nor was *P. stercorea* abundance (Low WAZ: r = +0.031, p = 0.627; High WAZ: r = −0.079, p = 0.414). These null associations indicate that differences in infection protection between strata are not attributable to variation in community diversity or colonisation level, and support a model in which WAZ primarily modifies host response capacity rather than microbiome composition.

### No association with weight gain

*P. stercorea* was not associated with weight gain (β = +0.028, 95% CI −0.017 to +0.073, p = 0.217), with <5% of this null association mediated by illness. There was no evidence of effect modification by WAZ (interaction p = 0.143; Supplementary Table S5; Supplementary Table S6), consistent with energy-allocation frameworks in which infection-mediated growth effects require concurrent nutritional constraint and support.

## Discussion

These findings resolve into two mechanistically distinct, compartment-specific pathways. Enteric outcomes are consistent with community-mediated colonisation resistance, whereas respiratory outcomes are richness-independent and attributable specifically to *P. stercorea*. Both pathways show host-context dependence: respiratory protection is strongest in Low-WAZ children - the same stratum in which Day 1 colonisation associates with suppressed CRP rather than elevated AGP - a co-localisation consistent with immune modulation as the operative mechanism rather than a non-specific effect of colonisation.

A key structural observation underpins this framework: WAZ is uncorrelated with microbiome richness and *P. stercorea* abundance within strata, arguing against models in which WAZ acts through community diversity, and instead supporting a role for host immune-metabolic reserve as a modifier of response capacity. In the respiratory compartment, the absence of attenuation after richness adjustment, together with null richness–ARI associations and significant WAZ interactions, supports a species-autonomous pathway whose expression depends on host immune context. In contrast, enteric outcomes show directionally consistent richness involvement: diarrhoea frequency showed 44% richness-mediated attenuation, with corroborating directionality from diarrhoea duration. Notably, the richness signal in enteric outcomes was restricted to Low-WAZ children; richness did not predict diarrhoea outcomes in High-WAZ children, suggesting that higher immune-metabolic reserve confers a colonisation resistance floor that attenuates the marginal contribution of *P. stercorea*-driven richness elevation. The same analytical framework thus discriminates between ecological mechanisms within a single population.

Temporally resolved biomarker patterns further refine this interpretation. Prospective D1→D1 associations indicate that *P. stercorea* colonisation in malnourished children is linked to reduced inflammatory burden, consistent with decreased pathogen-driven inflammation and/or altered innate immune set-points. In well-nourished children, the reverse D85→D1 pattern - baseline inflammation predicting subsequent colonisation - suggests that immune-active mucosal environments can sustain *P. stercorea* when metabolic resources are sufficient. Together, these findings define a nutrition-dependent, directionally asymmetric interaction between host inflammatory state and microbial colonisation.

The gut–lung axis provides a plausible, though untested, substrate for the respiratory pathway (Dang and Marsland, 2019; Wypych et al., 2019). Microbiome-derived metabolites and microbial ligands, including short-chain fatty acids and TLR2/TLR4 agonists, can prime innate immunity at distal mucosal sites such as the lung. The absence of a corresponding *P. copri* effect, despite co-occurrence and comparable contributions to richness, indicates that the operative signal is species-specific. Differences in metabolite profiles or cell-wall polysaccharide composition represent plausible mechanisms and are directly testable. The concentration of effects in ARI and fever - predominantly viral outcomes where colonisation resistance is unlikely to dominate, further supports systemic innate priming as a key pathway. In Low-WAZ children, a depressed inflammatory baseline may render the immune system more responsive to microbially-driven innate priming; in High-WAZ children, a pre-existing higher immune set-point may attenuate the marginal benefit of colonisation, consistent with the D85→D1 pattern in which inflammatory tone predicts colonisation rather than the reverse.

This species-resolved, compartmentalised framework refines ecological models of microbiome function. Protective effects are not reducible to diversity metrics alone but partition into community-level and species-autonomous components, with *P. stercorea* emerging as a candidate keystone taxon linking microbial ecology to host defence. Observed co-occurrence patterns are most parsimoniously explained by shared environmental selection rather than obligate functional interdependence, reinforcing attribution of the protective signal to *P. stercorea* itself.

These findings have direct implications for intervention design. Enteric protection appears to require community restoration, consistent with richness-dependent colonisation resistance and niche saturation. In contrast, respiratory protection is achievable through species-targeted approaches - a single-species probiotic or postbiotic strategy - without requiring wholesale restructuring of the gut community. The concentration of respiratory protection in hosts with low immune-metabolic reserve extends the translational logic to western populations: the low-richness, low-redundancy microbiome architecture characteristic of industrialised adults may represent a structurally analogous context to the high-protection paediatric phenotype rather than a categorical extrapolation from it. Nutritional status functions here as a modifier of host response capacity rather than a target for intervention - the implication for western markets is targeted microbiome support, not nutritional supplementation (Jin et al., 2021; GBD 2015 LRI Collaborators, 2017).

## Limitations

These analyses are observational. D85 as primary exposure may underestimate true associations through reverse causation, rendering estimates conservative (Supplementary Methods SM1). Specific causative pathogens were not characterised, precluding mechanistic resolution of the gut–lung axis hypothesis. The WAZ-stratified mediation analyses reveal directionally consistent patterns that invite prospective mechanistic testing in independent cohorts. The WAZ interaction test did not reach significance after BH correction across nine outcomes (ARI interaction q = 0.112), reflecting limited power in the High-WAZ stratum (n = 109), such that restriction of effect cannot be concluded; the stratum-specific betas are directionally consistent with host-context dependence but the data do not exclude meaningful protection in higher-immune-reserve hosts. Residual confounding cannot be excluded, though consistency across models and temporal ordering reduce this likelihood. The analyses were conducted in a single Gambian cohort; replication in independent paediatric populations with comparable microbiome profiling - such as PROVIDE (Bangladesh) (Kirkpatrick et al., 2015), WASH Benefits (Kenya/Bangladesh) (Null et al., 2018), or GEMS (Kotloff et al., 2013) - would strengthen generalisability. Within-cohort robustness is supported by temporal consistency of Path A estimates across D1, D15, and D85 and by treatment-arm sensitivity analyses (Supplementary Table S3).

In summary, *P. stercorea* is associated with reduced infection risk through two distinct ecological routes. Community-level restoration may address enteric outcomes; richness-independent, species-specific effects on respiratory immunity operate through a host-context-dependent pathway that is most active in low-immune-reserve hosts - a structural profile shared by nutritionally vulnerable children, western populations with reduced microbiome diversity and redundancy, the elderly, and individuals with chronic illness.

## Methods

### Ethics and cohort

This study uses data from the IHAT-GUT trial (NCT02941081), a three-arm randomised controlled trial conducted from November 2017 to November 2018 in rural communities of the Upper River Region, The Gambia. The study protocol was reviewed and approved by The Gambia Government/MRC Joint Ethics Committee (reference SCC1489), with Clinical Trials Authorisation granted by the Medicines Control Agency, The Gambia. The trial was conducted in accordance with the Declaration of Helsinki and ICH guidelines for Good Clinical Practice. Written informed consent was obtained from parents or legal guardians prior to enrolment. Full details of study design, participant eligibility, recruitment, microbiome profiling, and exclusion criteria are described in de Goffau et al., (2022) and Ofordile et al. (2025). Briefly, stool samples from 633 children aged 6–35 months with complete microbiome and adverse event data were included after excluding samples failing quality control and those from antibiotic-treated participants. As previously reported (Ofordile et al., 2025; de Goffau et al., 2022), the iron treatment arm had no significant effect on gut microbiome composition, alpha or beta diversity, or adverse event incidence, and its inclusion does not alter the conclusions of the present analyses (Supplementary Table S3). The treatment arm was therefore not included as a covariate in downstream models.

### Statistical analysis used in this study

#### Study data and preprocessing

Microbiome data and clinical metadata were derived from the IHAT-GUT clinical trial stool samples collected at Days 1 (D1), 15 (D15), and 85 (D85). The primary datasets used were *Merged_Illness_Cohorts_WithBiomarkers*.*csv*, containing microbiome taxonomic abundances and host metadata, and *Adverse Events*.*csv*, containing longitudinal illness records.

Species-level microbiome data (columns spanning all detected taxa) were aligned with clinical metadata using unique child identifiers. Relative abundances of *Prevotella stercorea* and *P. copri* were log-transformed (log_1+_x). Microbiome richness was defined as the number of taxa with non-zero abundance per sample.

Inflammation markers (CRP, AGP, calprotectin) were log-transformed to account for skewed distributions. Calprotectin values exceeding assay limits were truncated at 600 µg/g prior to transformation.

Adverse event (AE) data were summarised per participant as total frequency (number of episodes) and cumulative duration (days) for infection, diarrhoea, acute respiratory infection (ARI), and fever. With a follow-up period of 113 days, AE counts were capped (frequency ≤5; duration ≤30 days) to limit the influence of extreme outliers. ARI was defined as a clinician-diagnosed acute illness with cough and/or difficulty breathing, with or without fever, consistent with WHO/IMCI criteria for acute respiratory infection. ARI ascertainment encompassed both upper and lower respiratory tract presentations and was not stratified by anatomical site; specific causative pathogens (viral or bacterial) were not investigated.

ΔWeight was calculated as weight at D85 minus enrolment weight. Analyses were restricted to participants with complete covariates.

#### Community richness (Path A)

Associations between *Prevotella* abundance and microbiome richness were evaluated using linear regression models adjusted for age, sex, and baseline nutritional status (Supplementary Table S1):

lm(richness ∼ log_Pstercorea + age_sampling + gender_n + HAZ_enrolment, data = dat)

To integrate repeated measures across D1 and D15, linear mixed-effects models with random intercepts for participants were fitted:

lmer(richness ∼ log_Pstercorea + age_sampling + gender_n + HAZ_enrolment +

timepoint_num + (1 | rand_no), data = d1d15). All associations were highly significant across all timepoints (all p < 10^−6^); Benjamini–Hochberg correction was applied for completeness and q-values are reported alongside p-values in the table

#### Inflammation models (Path B1)

The relationship between *Prevotella* abundance and systemic inflammation biomarkers was assessed using linear models at individual timepoints (D1, D15) and combined mixed-effects models (Supplementary Table S4):

lm(log_CRP ∼ log_Pstercorea + HAZ_enrolment + age_sampling + gender_n, data = d1)

lmer(log_CRP ∼ log_Pstercorea + HAZ_enrolment + age_sampling + gender_n + timepoint_num + (1 | rand_no), data = d1d15)

Parallel models were fitted for calprotectin and AGP, and repeated for *P. copri* to assess species specificity. P-values from inflammation models were adjusted using the Benjamini–Hochberg procedure within each exposure block (P. stercorea and P. copri separately)

### Adverse event modelling

#### Count-based models (negative binomial GLMs)

Given overdispersion in AE frequency data, negative binomial generalized linear models were used (Supplementary Methods SM3):

glm.nb(ae_freq ∼ log_Pstercorea + age_enrolment + gender_n + HAZ_enrolment, data = dat)

To distinguish total versus richness-independent effects, richness-adjusted models were fitted:

glm.nb(ae_freq ∼ log_Pstercorea + richness + age_enrolment + gender_n + HAZ_enrolment, data = dat)

Species specificity was assessed using joint models including both taxa (Supplementary Methods SM4):

glm.nb(ae_freq ∼ log_Pstercorea + log_Pcopri + age_enrolment + gender_n + HAZ_enrolment, data = dat)

#### Illness and growth (Path B2)

The effect of infection burden on weight gain was evaluated using linear models (Supplementary Table S5):

lm(delta_Weight ∼ Ill_binary + HAZ_enrolment + age_enrolment + gender_n + Weight_enrolment, data = d85_growth)

Frequency- and duration-based AE metrics were analysed separately for infection, diarrhoea, fever, and acute respiratory infection (ARI). P-values from illness-to-weight models were adjusted using the Benjamini–Hochberg procedure across the nine illness-type comparisons

#### Causal chain and mediation framework (Path C)

To evaluate whether illness mediated the relationship between *Prevotella* abundance and growth, total and direct effects were estimated (Supplementary Methods SM2; Supplementary Table S6):

\# Total effect

lm(delta_Weight ∼ log_Pstercorea + covariates, data = d85_growth)

\# Direct effect (illness-adjusted)

lm(delta_Weight ∼ log_Pstercorea + Ill_binary + covariates, data = d85_growth)

The proportion mediated by illness was calculated as:

pct_mediated <-100 * (beta_total - beta_direct) / beta_total P-values from causal chain models (M1–M4) were adjusted using the Benjamini–Hochberg procedure within each model block

#### Effect modification by host immune context

Effect modification by nutritional status was assessed for the associations between *P. stercorea* abundance and (i) systemic inflammation biomarkers and (ii) weight gain. Analyses were conducted using enrolment z-scores (WAZ, HAZ, WHZ), dichotomised at the cohort median within each analytical dataset.

Two complementary approaches were applied (Supplementary Table S7). First, interaction models of the form:

lm(outcome ∼ log_Pstercorea * z_group + covariates, data = dat)

were fitted, where *z_group* is the median-split variable. The corresponding continuous z-score was excluded from covariates to avoid collinearity. The interaction term (*log_Pstercorea:z_groupHigh*) provided a formal test of effect modification, with the main effect representing the low z-score (reference) group.

Second, stratified models with identical covariate specifications were fitted within each stratum to obtain group-specific effect estimates.

Analyses were applied to:

- Inflammation models (M1): D85 *P. stercorea* → D1 biomarkers (CRP, calprotectin, AGP), adjusted for HAZ at enrolment, age at sampling, and sex
- Growth models (M4): D85 *P. stercorea* → ΔWeight, adjusted for HAZ at enrolment, age at enrolment, sex, and weight at enrolment

WAZ was treated as the stratification variable throughout.

To support mechanistic interpretation, additional diagnostic analyses were performed (Supplementary Table S8):

- Correlation structure: Pairwise Pearson correlations (r) between WAZ, microbiome richness, and *P. stercorea* abundance were assessed overall and within WAZ strata to exclude proxy relationships.
- Negative control analysis: Richness was evaluated as an alternative exposure in parallel models of the form *lm(outcome ∼ richness + covariates)*,
- Mediation analysis: Richness-mediated effects were quantified using a standard decomposition framework: *Total effect (c) = Direct effect (c′) + Indirect effect (a × b)*, models (M1): D85 *P. stercorea* → D1 biomarkers (CRP, calprotectin, AGP), adjusted for HAZ at enrolment, age at sampling, and sex

where *a* represents the association between *P. stercorea* and richness, and *b* the association between richness and outcome conditional on *P. stercorea*. Statistical inference was based on Sobel tests and nonparametric bootstrap resampling (mediation package, 1,000 simulations, seed 42), with percentage attenuation defined as *(c − c′)/c × 100. Sobel test p-values were adjusted using the Benjamini–Hochberg procedure within the P. stercorea exposure block; q-values are reported in table footnotes* to distinguish community-mediated from species-autonomous pathways. The primary pre-specified outcome was ARI frequency; P. copri comparisons addressed species specificity. P-values from negative binomial GLMs were adjusted for multiple comparisons using the Benjamini–Hochberg procedure (p.adjust, R), applied separately within the P. stercorea and P. copri exposure families; q-values are reported in table footnotes.

### Model outputs

Model estimates, confidence intervals, and p-values were extracted using the *broom* and *broom*.*mixed* packages, and all results were exported as structured tables:

tidy(fit, conf.int = TRUE)

write_xlsx(results_list, “Table_outputs.xlsx”)

## Supporting information

IHAT Paper2 Supplementary

## Data Availability

The datasets generated and/or analysed during the current study are available on Mendeley Data: https://data.mendeley.com/preview/fr5p4cgc7j?a=76a1e5f5-c508-4720-b445-45b75c0eee07. Raw bacterial 16S rRNA sequence data have been deposited in the European Nucleotide Archive (ENA) under accession number ERP110905.

## Software and Reproducibility

All analyses were conducted in R (v4.5.0; R Core Team) using *tidyverse, lme4, lmerTest, MASS, broom*, and *writexl*. Intermediate .rds objects were employed to standardize datasets across analytical steps and ensure full reproducibility. Datasets were prepared in Microsoft Excel (2016; Microsoft Corporation, Redmond, WA, USA) and analyzed in R within RStudio (v2025.09.1, ‘Cucumberleaf Sunflower’; Posit Software, PBC) using Quarto (v1.7.32) on Windows 10 for reproducible reporting and figure generation. Figure panels were assembled and formatted in Inkscape (v1.4.2). The complete reproducible analysis pipeline is available at https://github.com/ofordile-star/IHAT_Pstercorea and will be publicly archived upon submission.

## Acknowledgements

I acknowledge the staff of the Medical Research Council (MRC) Unit, The Gambia at the London School of Hygiene and Tropical Medicine (LSHTM), in particular Mr. Bakary Baldeh, the IHAT-GUT Nurse Coordinator, and the team of study nurses for their meticulous recording and management of adverse events and care of study participants. I also thank the dedicated IHAT-GUT field team for their invaluable support and contribution to this research. I am grateful to the staff of Yorrobawol Health Centre, Darsilami Community Health Post, Konkuba Community Health Post, Taibatu Health Post, and Chamoi Health Centre for welcoming our team and supporting our study. I sincerely thank the local communities, study participants, and their families in the Wuli and Sandu districts of the Upper River Region, The Gambia, for their generous participation in this study.

The IHAT-GUT study was supported by a Bill & Melinda Gates Foundation Grand Challenges New Interventions in Global Health award (OPP1140952). The Nutrition Group of the MRC Unit The Gambia at LSHTM is supported by core funding (MC-A760-5QX00) to the MRC Unit The Gambia/MRC International Nutrition Group by the UK Medical Research Council (MRC) and the UK Department for International Development (DFID) under the MRC/DFID Concordat agreement. The funders had no role in study design, data collection and analysis, decision to publish, or preparation of the manuscript.

## Author Contributions

O.O. oversaw adverse event data collection, conceived the study, established the study design, developed the analytical framework, conducted all analyses, interpreted the results, and wrote the manuscript.

## References

1. Chen RY, et al. 2021. A microbiota-directed food intervention for undernourished children. New England Journal of Medicine 384:1517–1528. doi:10.1056/NEJMoa2023294

2. Dang AT, Marsland BJ. 2019. Microbes, metabolites, and the gut–lung axis. Mucosal Immunology 12:843–850. doi:10.1038/s41385-019-0160-6

3. De Filippo C, Cavalieri D, Di Paola M, Ramazzotti M, Poullet JB, Massart S, Collini S, Pieraccini G, Lionetti P. 2010. Impact of diet in shaping gut microbiota revealed by a comparative study in children from Europe and rural Africa. Proceedings of the National Academy of Sciences 107:14691–14696. doi:10.1073/pnas.1005963107

4. de Goffau MC, Jallow AT, Sanyang C, Prentice AM, Meagher N, Price DJ, Revill PA, Parkhill J, Pereira DIA, Wagner J. 2022. Gut microbiomes from Gambian infants reveal the development of a non-industrialized Prevotella-based trophic network. Nature Microbiology 7:132–144. doi:10.1038/s41564-021-01023-6

5. Foster KR, Bell T. 2012. Competition, not cooperation, dominates interactions among culturable microbial species. Current Biology 22:1845–1850. doi:10.1016/j.cub.2012.08.005

6. GBD 2015 LRI Collaborators. 2017. Estimates of the global, regional, and national morbidity, mortality, and aetiologies of lower respiratory tract infections in 195 countries: a systematic analysis for the Global Burden of Disease Study 2015. Lancet Infectious Diseases 17:1133–1161. doi:10.1016/S1473-3099(17)30396-1

7. GBD 2019 Diseases and Injuries Collaborators. 2020. Global burden of 369 diseases and injuries in 204 countries and territories, 1990–2019: a systematic analysis for the Global Burden of Disease Study 2019. Lancet 396:1204–1222. doi:10.1016/S0140-6736(20)30925-9

8. Jin X, Ren J, Li R, Gao Y, Zhang H, Li J, Zhang J, Wang X, Wang G. 2021. Global burden of upper respiratory infections in 204 countries and territories, from 1990 to 2019. EClinicalMedicine 37:100986. doi:10.1016/j.eclinm.2021.100986

9. Kirkpatrick BD, et al. 2015. The “Performance of Rotavirus and Oral Polio Vaccines in Developing Countries” (PROVIDE) Study: description of methods of an interventional study designed to explore complex biologic problems. American Journal of Tropical Medicine and Hygiene 92:744–751. doi:10.4269/ajtmh.14-0518

10. Kotloff KL, et al. 2013. Burden and aetiology of diarrhoeal disease in infants and young children in developing countries (the Global Enteric Multicenter Study, GEMS): a prospective, case-control study. Lancet 382:209–222. doi:10.1016/S0140-6736(13)60844-2

11. Mostafa I, Hibberd MC, Hartman SJ, Rahman MDHH, Mahfuz M, Hasan SMT, Ashorn P, Barratt MJ, et al. 2024. A microbiota-directed complementary food intervention in 12–18-month-old Bangladeshi children improves linear growth. EBioMedicine 104:105166. doi:10.1016/j.ebiom.2024.105166

12. Null C, et al. 2018. Effects of water quality, sanitation, handwashing, and nutritional interventions on diarrhoea and child growth in rural Kenya: a cluster-randomised controlled trial. Lancet Global Health 6:e316–e329. doi:10.1016/S2214-109X(18)30005-6

13. Ofordile O, Pereira DIA, Prentice AM, Wagner J. 2025. Gut Prevotella stercorea associates with protection against infection in rural African children. Nature Communications 16:11101. doi:10.1038/s41467-025-66011-4

14. Smits SA, Leach J, Sonnenburg ED, Gonzalez CG, Lichtman JS, Reid G, Knight R, Sonnenburg JL. 2017. Seasonal cycling in the gut microbiome of the Hadza hunter-gatherers of Tanzania. Science 357:802–806. doi:10.1126/science.aan4834

15. Spragge F, Bansept F, Thöle D, Wulff T, Quax TEF, Buckling A, Friman VP, Kümmerli R, De Monte S, Foster KR. 2023. Microbiome diversity protects against pathogens by nutrient blocking. Science 382:eadj3502. doi:10.1126/science.adj3502

16. Tickell KD, Brander RL, Walson JL, Denno DM. 2020. The effect of acute malnutrition on enteric pathogens, moderate-to-severe diarrhoea, and associated mortality in the Global Enteric Multicenter Study cohort: a post-hoc analysis. Lancet Global Health 8:e215–e224. doi:10.1016/S2214-109X(19)30484-X

17. Wypych TP, Wickramasinghe LC, Marsland BJ. 2019. The influence of the microbiome on respiratory health. Nature Immunology 20:1279–1290. doi:10.1038/s41590-019-0451-9

