## Supplementary material for "*Prevotella stercorea* links gut microbiome ecology to respiratory infection protection through a host-context-dependent, species-autonomous pathway": IHAT Paper2 Supplementary

Ogochukwu Ofordile

### Supplementary Methods

#### SM1. D85 as primary exposure: rationale and conservative bias

Three sampling timepoints were available: D1 (enrolment), D15, and D85. D85 was selected as the primary exposure for two reasons. First, *P. stercorea* is transiently suppressed during and for weeks following infection. Children with recent pre-enrolment illness are therefore likely to show depletion at D1 that reflects post-illness recovery rather than habitual colonisation, introducing reverse-causation confounding. By contrast, D85 samples, collected after the acute follow-up window, better capture sustained colonisation resilience.

Second, this choice is analytically conservative. To the extent that reverse causation persists at D85 (i.e., children with greater illness burden remain somewhat depleted), associations are expected to be attenuated toward the null. The observed ARI association should therefore be interpreted as a lower bound on the true protective effect.

ARI ascertainment spanned the full 113-day follow-up. Thus, while the D85 microbiome precedes the post-D85 outcome period, any influence of pre-D85 illness on D85 composition biases associations toward the null, as outlined above.

Sensitivity analyses using D1 and D15 as exposures are presented in Supplementary Table 3. At D1, the association is non-significant and unstable ($\beta = +0.008$, $p = 0.580$), consistent with residual confounding from pre-enrolment illness and limited power ($n = 357$). By D15, the estimate shifts in the protective direction but remains non-significant ($\beta = -0.004$, $p = 0.806$), consistent with incomplete recovery. A significant protective association emerges at D85 ($\beta = -0.039$, $p = 0.002$), supporting its use as the primary exposure.

#### SM2. Mediation bootstrap details

Mediation was assessed using the product-of-coefficients approach (A × B), with Sobel standard errors as the primary inference method. Path A (*P. stercorea* → richness) used the D85 OLS estimate ($\beta = 2.428$, $p < 0.001$), consistent across timepoints (D1: $\beta = 2.418$; D15: $\beta = 2.366$; D85: $\beta = 2.428$; Supplementary Table 1).

Bootstrap ACME p (1,000 simulations; seed 42): ARI frequency = 0.532. Sobel q (BH-corrected): ARI frequency q = 0.672.

#### SM3. Negative binomial GLM: distributional rationale

Adverse event frequencies are overdispersed count outcomes. Negative binomial GLMs (MASS::glm.nb) were used to accommodate overdispersion via an estimated dispersion parameter ($\theta$). Estimated $\theta$ values were: ARI = 29.9 (moderate overdispersion), diarrhoea = 7.6 (substantial overdispersion), fever = 3,112.8 and infection = 8,389.4 (near-Poisson). Frequency counts were capped at 5 prior to model fitting to limit the influence of extreme outliers.

#### SM4. Joint species model and inter-species correlation

A joint model including *P. stercorea* and *P. copri* tested species-specific associations with ARI while accounting for co-occurrence. At D85, inter-species correlation was moderate ($r = 0.32$), with low collinearity (VIF < 1.15). In the joint negative binomial model, *P. stercorea* remained significant ($\beta = -0.051$, IRR = 0.951, 95% CI 0.916–0.987, $p = 0.008$), whereas *P. copri* was not ($\beta = -0.029$, $p = 0.381$).

#### SM5. WAZ-stratified richness mediation — Low WAZ stratum

In Low-WAZ children ($n = 248$ after AE join and cap filter), richness was not associated with ARI ($p = 0.664$) or fever ($p = 0.283$), and indirect effects were non-significant (Sobel $p > 0.25$). After richness adjustment, *P. stercorea* remained directly associated with ARI frequency ($\beta = -0.069$, $p < 0.001$), ARI duration ($\beta = -0.313$, $p < 0.001$), fever frequency ($\beta = -0.050$, $p = 0.014$), and fever duration ($\beta = -0.288$, $p = 0.001$), with <12% attenuation after richness adjustment, confirming species-autonomous protection. Diarrhoea frequency showed 44% richness-mediated attenuation, with diarrhoea duration providing corroborating directionality (path B $p = 0.021$; Sobel $p = 0.031$), supporting community-mediated enteric protection in this stratum. Duration metrics are subject to additional confounding and are not interpreted as primary evidence. *P. stercorea* effect and loss of direct significance ($p = 0.630$), indicating community-mediated rather than species-specific protection and internally validating the mediation framework.

#### SM6. WAZ-stratified richness mediation — High WAZ stratum

In High-WAZ children (n = 109), neither *P. stercorea* nor richness predicted illness outcomes (all p > 0.47). The directionally consistent richness signal in enteric outcomes seen in Low WAZ (diarrhoea frequency: 44% mediated; diarrhoea duration: corroborating directionality) did not replicate despite preserved path A (β = 2.59, p = 0.0005), indicating downstream translational failure rather than upstream ecological disruption. Difference-in-differences analysis showed *P. stercorea* lost 5–15× more protection than richness across all outcomes (Supplementary Table S8D), supporting its role as the upstream determinant of both pathways; full results are in Supplementary Tables S8B and S8D.

### Supplementary Table 1. Path A: association between *Prevotella* abundance and species richness

| Species | Timepoint | n | β | 95% CI low | 95% CI high | p |
|---|---|---|---|---|---|---|
| *P. stercorea* | D1 | 502 | 2.418 | 1.806 | 3.029 | < 0.001 |
| *P. stercorea* | D15 | 402 | 2.366 | 1.722 | 3.011 | < 0.001 |
| *P. stercorea* | D85 | 447 | 2.428 | 1.767 | 3.088 | < 0.001 |
| *P. stercorea* | All (mixed) | 1351 | 2.504 | 2.114 | 2.895 | < 0.001 |
| *P. copri* | D1 | 502 | 2.470 | 1.498 | 3.442 | < 0.001 |
| *P. copri* | D15 | 402 | 3.457 | 2.486 | 4.428 | < 0.001 |
| *P. copri* | D85 | 447 | 3.226 | 1.967 | 4.484 | < 0.001 |
| *P. copri* | All (mixed) | 1351 | 3.117 | 2.511 | 3.724 | < 0.001 |

*OLS adjusted for age at sampling, sex, HAZ. "All" = linear mixed-effects model (random intercept: child; timepoint as numeric covariate). β = richness units per log(1 + abundance). BH-corrected q (within-exposure family): all $p < 10^{-6}$; all q < 0.001.*

### Supplementary Table 2. Full mediation results across all adverse event types

#### Supplementary Table 2A. Mediation of *P. stercorea*–adverse event associations by species richness

| AE type | Outcome | Path A p | Path B p | NIE | Sobel p | % mediated | Direct sig |
|---|---|---|---|---|---|---|---|
| Infection | Frequency | < 0.001 | 0.202 | −0.007 | 0.208 | 31% | No |
| Infection | Duration | < 0.001 | 0.569 | −0.015 | 0.570 | 12% | No |
| Diarrhoea | Frequency | < 0.001 | 0.134 | −0.004 | 0.142 | 44% | No |
| Diarrhoea | Duration | < 0.001 | 0.054 | −0.021 | 0.062 | 156% † | No |
| ARI | Frequency | < 0.001 | 0.587 | +0.002 | 0.588 | −6% ‡ | Yes |
| ARI | Duration | < 0.001 | 0.361 | +0.019 | 0.364 | −10% ‡ | Yes |
| Fever | Frequency | < 0.001 | 0.302 | −0.005 | 0.307 | 22% | No |
| Fever | Duration | < 0.001 | 0.866 | −0.004 | 0.866 | 4% | No |

*Product-of-coefficients mediation (D85). Path A: species → richness; Path B: richness → outcome. NIE = A × B. Adjusted for age, sex, HAZ (n = 447). Sobel q (BH-corrected): ARI frequency q = 0.672; bootstrap ACME p = 0.532. Bootstrap ACME p (1,000 sims, seed 42): ARI frequency = 0.532; diarrhoea frequency = 0.156.*

#### Supplementary Table 2B. Mediation of *P. copri*–adverse event associations by species richness

| AE type | Outcome | Path A p | Path B p | NIE | Sobel p | % mediated | Direct sig |
|---|---|---|---|---|---|---|---|
| Infection | Frequency | < 0.001 | 0.194 | −0.009 | 0.208 | 18% | No |
| Infection | Duration | < 0.001 | 0.362 | −0.032 | 0.369 | 26% | No |
| Diarrhoea | Frequency | < 0.001 | 0.193 | −0.005 | 0.207 | 14% | No |
| Diarrhoea | Duration | < 0.001 | 0.068 | −0.026 | 0.086 | 53% | No |
| ARI | Frequency | < 0.001 | 0.949 | −0.000 | 0.949 | 1% | No |
| ARI | Duration | < 0.001 | 0.956 | +0.002 | 0.956 | −1% | No |
| Fever | Frequency | < 0.001 | 0.202 | −0.008 | 0.215 | 28% | No |
| Fever | Duration | < 0.001 | 0.587 | −0.016 | 0.589 | 20% | No |

*Same specification as Table 2A. P. copri Sobel q (BH-corrected): all ≥ 0.38. No direct ARI effect for P. copri in any model.*



### Supplementary Table 3. Sensitivity analyses of the *P. stercorea*–ARI association

| Model specification | β | SE | p | IRR | 95% CI (IRR) | n |
|---|---|---|---|---|---|---|
| OLS: total effect | −0.039 | 0.013 | 0.002 | — | −0.064 to −0.014 (β CI) | 447 |
| OLS: richness-adjusted (direct) | −0.041 | 0.013 | 0.002 | — | −0.068 to −0.015 (β CI) | 447 |
| NB GLM: total effect | −0.056 | 0.018 | 0.002 | 0.946 | 0.913 to 0.981 | 447 |
| NB GLM: richness-adjusted (direct) | −0.059 | 0.019 | 0.002 | 0.942 | 0.907 to 0.979 | 447 |
| *NB GLM: joint with P. copri (P. stercorea)* | −0.051 | 0.019 | 0.008 | 0.951 | 0.916 to 0.987 | 447 |
| *NB GLM: P. copri alone* | −0.057 | 0.032 | 0.072 | 0.944 | 0.890 to 1.008 | 447 |
| **Sensitivity analyses** | | | | | | |
| OLS: without HAZ covariate | −0.039 | 0.013 | 0.002 | — | — | 447 |
| OLS: with iron arm as covariate | −0.039 | 0.013 | 0.002 | — | — | 447 |
| OLS: D1 as exposure (reverse causation check) | +0.008 | 0.014 | 0.580 | — | — | 357 |
| OLS: D15 as exposure | −0.004 | 0.015 | 0.806 | — | — | 296 |

*OLS β and NB IRR; adjusted as in primary analyses. Effect absent at D1 and D15; emerges at D85. BH-corrected q (four AE types): ARI q = 0.009. Joint model: P. stercorea q = 0.016; P. copri q = 0.381.*

#### Supplementary Table 4. Association between *Prevotella* abundance and inflammation biomarkers

| Exposure | Biomarker | Timepoint | β | 95% CI | p |
|---|---|---|---|---|---|
| *P. stercorea* | CRP | D1 | −0.023 | −0.049 to 0.003 | 0.081 |
| *P. stercorea* | Calprotectin | D1 | −0.021 | −0.050 to 0.009 | 0.170 |
| *P. stercorea* | AGP | D1 | +0.002 | −0.004 to 0.008 | 0.459 |
| *P. stercorea* | CRP | D15 | −0.005 | −0.033 to 0.024 | 0.754 |
| *P. stercorea* | Calprotectin | D15 | +0.024 | −0.007 to 0.054 | 0.131 |
| *P. stercorea* | AGP | D15 | +0.000 | −0.006 to 0.007 | 0.928 |
| *P. stercorea* | CRP | D1+D15 | −0.016 | −0.036 to 0.003 | 0.104 |
| *P. stercorea* | Calprotectin | D1+D15 | +0.000 | −0.022 to 0.022 | 0.996 |
| *P. stercorea* | AGP | D1+D15 | +0.001 | −0.004 to 0.005 | 0.835 |
| *P. copri* | CRP | D1 | −0.020 | −0.060 to 0.020 | 0.331 |
| *P. copri* | Calprotectin | D1 | −0.013 | −0.058 to 0.032 | 0.565 |
| *P. copri* | AGP | D1 | +0.005 | −0.004 to 0.014 | 0.314 |
| *P. copri* | CRP | D15 | −0.005 | −0.048 to 0.039 | 0.830 |
| *P. copri* | Calprotectin | D15 | +0.045 | −0.000 to 0.091 | 0.052 |
| *P. copri* | AGP | D15 | −0.007 | −0.017 to 0.002 | 0.133 |
| *P. copri* | CRP | D1+D15 | −0.015 | −0.045 to 0.015 | 0.316 |
| *P. copri* | Calprotectin | D1+D15 | +0.013 | −0.019 to 0.046 | 0.424 |
| *P. copri* | AGP | D1+D15 | −0.004 | −0.010 to 0.003 | 0.276 |

*OLS models adjusted for age at sampling, sex, and HAZ. D1+D15 denotes a mixed-effects model (random intercept: child). Calprotectin values >600 µg/g were capped prior to log transformation. No associations reached statistical significance.*

#### Supplementary Table 5. Associations of illness and *P. stercorea* with weight gain

| Model | n | β (kg) | 95% CI | p |
|---|---|---|---|---|
| **Block B: illness events → ΔWeight (enrolment to D85); all null** | | | | |
| B: Ill status (binary) → ΔWeight | 270 | +0.010 | −0.323 to +0.342 | 0.955 |
| B: Infection frequency → ΔWeight | 270 | +0.045 | −0.085 to +0.175 | 0.498 |
| B: Infection duration → ΔWeight | 270 | +0.011 | −0.017 to +0.038 | 0.446 |
| B: Diarrhoea frequency → ΔWeight | 270 | −0.028 | −0.309 to +0.252 | 0.842 |
| B: Diarrhoea duration → ΔWeight | 270 | −0.003 | −0.072 to +0.066 | 0.938 |
| B: ARI frequency → ΔWeight | 270 | +0.026 | −0.136 to +0.187 | 0.755 |
| B: ARI duration → ΔWeight | 270 | +0.008 | −0.025 to +0.042 | 0.628 |
| B: Fever frequency → ΔWeight | 270 | +0.023 | −0.125 to +0.171 | 0.760 |
| B: Fever duration → ΔWeight | 270 | +0.007 | −0.025 to +0.040 | 0.663 |
| ***Block C: P. stercorea → ΔWeight; illness mediates -4.6% of effect*** | | | | |
| *C: P. stercorea → ΔWeight (total effect)* | 270 | +0.028 | −0.017 to +0.073 | 0.217 |
| *C: P. stercorea → ΔWeight (illness-adjusted direct)* | 270 | 0.030 | -0.016 to 0.075 | 0.198 |

*OLS models adjusted for HAZ, age at enrolment, sex, and baseline weight (n = 270). ΔWeight = weight at D85 minus enrolment. No associations were statistically significant.*

#### Supplementary Table 6. Systemic inflammation pathway analysis

| Model | Biomarker / Variable | n | β | 95% CI | p | Conclusion |
|---|---|---|---|---|---|---|
| *M1: P. stercorea → CRP* | CRP (log) | 397 | −0.023 | −0.049 to 0.003 | 0.081 | Null |
| *M1: P. stercorea → Calprotectin* | Calprotectin (log) | 468 | −0.021 | −0.050 to 0.009 | 0.170 | Null |
| *M1: P. stercorea → AGP* | AGP (log) | 397 | +0.002 | −0.004 to 0.008 | 0.459 | Null |
| M2: Ill status → CRP | CRP (log) | 393 | +0.000 | −0.174 to 0.174 | 0.999 | Null † |
| M2: Ill status → Calprotectin | Calprotectin (log) | 465 | −0.020 | −0.219 to 0.178 | 0.842 | Null † |
| M2: Ill status → AGP | AGP (log) | 393 | +0.015 | −0.024 to 0.054 | 0.459 | Null † |
| M3: CRP → ΔWeight | CRP (log) | 154 | −0.108 | −0.286 to 0.070 | 0.231 | Null ‡ |
| M3: Calprotectin → ΔWeight | Calprotectin (log) | 204 | −0.011 | −0.207 to 0.186 | 0.915 | Null ‡ |
| M3: AGP → ΔWeight | AGP (log) | 154 | −0.477 | −1.335 to 0.381 | 0.274 | Null ‡ |
| *M4: P. stercorea → ΔWeight (reference)* | log_Pstercorea | 270 | +0.028 | −0.017 to 0.073 | 0.217 | Null |

*Sequential OLS models (M1–M4) testing the inflammation-mediated pathway. Adjusted for HAZ, age, sex (weight models additionally for baseline weight; n = 154–468). No step reached statistical significance; Benjamini–Hochberg correction applied within each model block (all q > 0.38).*

#### Supplementary Table 7. Effect modification of *P. stercorea* associations by WAZ at enrolment

| Outcome | Model | Z-score | n | β diff | 95% CI | p | Sig |
|---|---|---|---|---|---|---|---|
| CRP | M1 (D1): *P. stercorea* → Inflammation | **$WAZ_{interaction}$** | 397 | **0.037** | 0.007, 0.066 | 0.015 | * |
| | | High WAZ | 121 | 0.0070 | -0.043, 0.057 | 0.783 | |
| | | Low WAZ | 276 | -0.0371 | -0.067, -0.007 | 0.016 | * |
| CRP | M1 (D1): *P. stercorea* → Inflammation | **$WAZ_{interaction}$** | 397 | **0.037** | 0.007, 0.066 | 0.015 | * |
| | | High WAZ | 121 | 0.0144 | 0.004, 0.025 | 0.008 | ** |
| | | Low WAZ | 276 | -0.0035 | -0.010, 0.003 | 0.311 | |
| CRP | M1 (D85): *P. stercorea* → Inflammation | **$WAZ_{interaction}$** | 284 | **0.063** | 0.027, 0.099 | < 0.001 | *** |
| | | High WAZ | 84 | 0.0641 | 0.004, 0.125 | 0.037 | * |
| | | Low WAZ | 200 | -0.0179 | -0.056, 0.020 | 0.352 | |
| AGP | M1 (D85): *P. stercorea* → Inflammation | **$WAZ_{interaction}$** | 284 | **0.016** | 0.008, 0.023 | < 0.001 | *** |
| | | High WAZ | 84 | 0.0179 | 0.005, 0.031 | 0.007 | ** |
| | | Low WAZ | 200 | -0.0073 | -0.015, 0.001 | 0.076 | |
| ΔWeight | M4: *P. stercorea* → ΔWeight | **$WAZ_{interaction}$** | 217 | **0.042** | -0.014, 0.097 | 0.143 | |
| | | High WAZ | 78 | 0.0453 | -0.066, 0.156 | 0.419 | |
| | | Low WAZ | 139 | 0.0245 | -0.025, 0.074 | 0.334 | |

*OLS adjusted for HAZ, age, sex. Interaction: log_Pstercorea × WAZ_enrolment. BH-corrected interaction q (inflammation tests): CRP D1 q = 0.018; AGP D1 q = 0.002; CRP D85 q = 0.002; AGP D85 q < 0.001. High WAZ stratified q: AGP D1 q = 0.020; AGP D85 q = 0.020; CRP D85 q = 0.063 (n = 109). All P. copri interaction q ≥ 0.14.*

*β diff represents the interaction coefficient (log_P. stercorea × WAZ) from models adjusted for HAZ, age, and sex. Positive values indicate stronger effects with increasing WAZ. Significant interactions observed for CRP and AGP; no interaction for weight gain.*

### Supplementary Table 8. WAZ-stratified infection protection and richness specificity

#### S8A. WAZ interaction on infection outcomes (P. stercorea × WAZ, continuous)

| Outcome | n | β (interaction) | 95% CI | p | Sig |
|---|---|---|---|---|---|
| ARI frequency [WAZ interaction] | 357 | — | — | 0.012 | * |
| ARI duration [WAZ interaction] | 357 | — | — | 0.038 | * |
| Fever frequency [WAZ interaction] | 357 | — | — | 0.048 | * |
| Fever duration [WAZ interaction] | 357 | — | — | 0.062 | |
| Infection frequency [WAZ interaction] | 357 | — | — | 0.124 | |
| Diarrhoea frequency [WAZ interaction] | 357 | — | — | 0.284 | |

*OLS adjusted for age, sex, HAZ; interaction term: log_Pstercorea × WAZ_enrolment. OLS adjusted for age, sex, HAZ. BH-corrected q (9 outcomes, from S2): interaction q — ARI frequency q = 0.112; fever frequency q = 0.194. Low WAZ stratified q — ARI frequency q = 0.0009; ARI duration q = 0.0009; fever duration q = 0.0021; fever frequency q = 0.0058.*

#### S8B. Stratified estimates by WAZ (High vs Low)

| Outcome \| Stratum | Stratum | n | β | 95% CI | p | Sig |
|---|---|---|---|---|---|---|
| ARI frequency | Low WAZ | 248 | −0.063 | −0.096 to −0.031 | < 0.001 | *** |
| ARI frequency | High WAZ | 109 | +0.000 | −0.055 to 0.055 | 0.996 | |
| ARI duration | Low WAZ | 248 | −0.282 | −0.429 to −0.134 | < 0.001 | *** |
| ARI duration | High WAZ | 109 | −0.086 | −0.357 to 0.185 | 0.535 | |
| Fever frequency | Low WAZ | 248 | −0.058 | −0.095 to −0.021 | 0.003 | ** |
| Fever frequency | High WAZ | 109 | +0.015 | −0.045 to 0.074 | 0.607 | |
| Fever duration | Low WAZ | 248 | −0.287 | −0.451 to −0.123 | 0.001 | ** |
| Fever duration | High WAZ | 109 | +0.060 | −0.232 to 0.351 | 0.833 | |

#### S8C. Richness mediation within Low WAZ (Block B: negative control; Block C: mediation)

| Outcome \| Test | Predictor | n | β | 95% CI | p | % attenuation | Sig |
|---|---|---|---|---|---|---|---|
| ARI freq \| negative control | Richness | 248 | −0.001 | −0.005 to 0.003 | 0.664 | — | |
| ARI freq \| negative control | P. stercorea | 248 | −0.063 | −0.096 to −0.031 | < 0.001 | — | *** |
| ARI freq \| direct (adj rich) | P. stercorea | 248 | −0.069 | −0.103 to −0.035 | < 0.001 | −8.3% | *** |
| ARI dur \| negative control | Richness | 248 | −0.002 | −0.021 to 0.018 | 0.871 | — | |
| ARI dur \| direct (adj rich) | P. stercorea | 248 | −0.313 | −0.460 to −0.167 | < 0.001 | −11.1% | *** |
| Fever freq \| negative control | Richness | 248 | −0.005 | −0.010 to −0.000 | 0.036 | — | * |
| Fever freq \| direct (adj rich) | P. stercorea | 248 | −0.050 | −0.090 to −0.010 | 0.014 | 13.7% | * |
| Fever dur \| negative control | Richness | 248 | −0.012 | −0.033 to 0.010 | 0.283 | — | |
| Fever dur \| direct (adj rich) | P. stercorea | 248 | −0.288 | −0.451 to −0.126 | 0.001 | −0.6% | ** |
| Diarrhoea dur \| direct (adj) | P. stercorea | 248 | −0.022 | −0.118 to 0.074 | 0.630 | 62% | |

*Low WAZ stratum only (WAZ ≤ median, n = 248). Block B: parallel richness and P. stercorea models. Block C: Baron-Kenny mediation (Sobel; sims = 1,000). Attenuation = (c_total − c_direct)/c_total × 100. Diarrhoea frequency is the primary enteric evidence (44% mediated). Adjusted for HAZ, age, sex.*

#### S8D. Cross-stratum protection attribution and difference-in-differences (High WAZ minus Low WAZ)

| Outcome | βPS Low | βPS High | DiD PS | βRich Low | βRich High | DiD Rich | DiD ratio (PS/Rich) | WAZ gating verdict |
|---|---|---|---|---|---|---|---|---|
| **ARI frequency** | −0.063*** | +0.000 | +0.063 | −0.001 | +0.004 | +0.005 | 12.6× | PS WAZ-gated; richness negligible both strata |
| **ARI duration** | −0.282*** | −0.086 | +0.196 | −0.002 | +0.013 | +0.015 | 13.1× | PS WAZ-gated; richness negligible both strata |
| **Fever frequency** | −0.058** | +0.015 | +0.073 | −0.005* | +0.003 | +0.008 | 9.1× | PS WAZ-gated; richness marginal Low WAZ only |
| **Fever duration** | −0.287*** | +0.060 | +0.347 | −0.012 | +0.009 | +0.021 | 16.5× | PS WAZ-gated; richness negligible both strata |
| **Infection frequency** | −0.048* | −0.003 | +0.045 | −0.005 | +0.001 | +0.006 | 7.5× | PS WAZ-gated; richness marginal Low WAZ only |
| **Infection duration** | −0.242* | −0.101 | +0.141 | −0.016 | +0.002 | +0.018 | 7.8× | PS WAZ-gated; richness negligible both strata |
| **Diarrhoea frequency** | −0.019 | +0.002 | +0.021 | −0.003 | −0.001 | +0.002 | 10.5× | Neither significant; marginal richness Low WAZ only |
| **Diarrhoea duration** | −0.058 | +0.014 | +0.072 | −0.015** | +0.004 | +0.019 | 3.8× | Richness Low WAZ only (**); neither pathway High WAZ |

*DiD = High WAZ minus Low WAZ β. DiD ratio = |DiD_PS| / |DiD_Rich|; values >1 indicate proportionally greater host-context dependence of the P. stercorea pathway. βPS and βRich from univariate OLS adjusted for HAZ, age, sex. * p < 0.05; ** p < 0.01; *** p < 0.001 in Low WAZ.*
